# Alignment-free metal ion-binding site prediction from protein sequence through pretrained language model and multi-task learning

**DOI:** 10.1101/2022.05.20.492769

**Authors:** Qianmu Yuan, Sheng Chen, Yu Wang, Huiying Zhao, Yuedong Yang

## Abstract

More than one-third of the proteins contain metal ions in the Protein Data Bank. Correct identification of metal ion-binding residues is important for understanding protein functions and designing novel drugs. Due to the small size and high versatility of metal ions, it remains challenging to computationally predict their binding sites from protein sequence. Existing sequence-based methods are of low accuracy due to the lack of structural information, and time-consuming owing to the usage of multi-sequence alignment. Here, we propose LMetalSite, an alignment-free sequence-based predictor for binding sites of the four most frequently seen metal ions (Zn^2+^, Ca^2+^, Mg^2+^ and Mn^2+^). LMetalSite leverages the pretrained language model to rapidly generate informative sequence representations and employs transformer to capture long-range dependencies. Multi-task learning is adopted to compensate for the scarcity of training data and capture the intrinsic similarities between different metal ions. LMetalSite was shown to surpass state-of-the-art structure-based methods by more than 19.7%, 14.4%, 36.8%, and 12.6% in AUPR on the four independent tests, respectively. Further analyses indicated that the self-attention modules are effective to learn the structural contexts of residues from protein sequence.

## 1. Introduction

Almost 40% of the proteins in the Protein Data Bank (PDB) [1] bind to metal ions [2], which are indispensable for the protein structural stability [3] and biological functions in cells such as enzyme catalysis and regulation of gene expression [4-6]. For example, Zn^2+^ ions can bind with specific nucleases and transcription factors to form Zn finger domains that recognize DNA and RNA for regulation of gene expression [6]. Hence, identifying amino acids involved in protein-metal-ion interactions helps to understand protein functions and design novel drugs [7]. Unfortunately, experimental methods for metal ion-binding site detection such as nuclear magnetic resonance [8] and absorption spectroscopy [9] are costly and time-consuming. Therefore, it is desirable to develop computational methods for making reliable metal ion-binding site prediction.

Many computational methods have been developed for predicting metal ion-binding sites, but the problem remains challenging due to the small size and high versatility of metal ions. Current methods can be classified into structure-based and sequence-based methods according to their used information. Structure-based approaches using experimental structures as input are often more accurate, which can be generally categorized into template-based methods, machine-learning-based methods, and hybrid methods. Template-based methods such as MIB [10] employ alignment algorithms to transfer the structure information of templates for binding site inference. Nevertheless, these methods will be seriously restricted when no high-quality template can be found. Structure-based machine learning methods handle protein structures by extracting geometric features and then feeding to the neural networks, or explicitly taking the structural context topology into account and training in an end-to-end way. One example of the former is DELIA [11], which treats protein structures as 2D images and uses convolutional neural networks to extract characteristics from protein distance matrices. Alternately, an example of the latter is GraphBind [12], which encodes protein structures as graphs and adopts graph neural networks to learn the local tertiary patterns for binding site prediction. Hybrid methods such as COACH [7] and IonCom [13] integrate template-based methods and machine-learning-based methods simultaneously. Albeit powerful, the structure-based methods are not applicable to most proteins whose tertiary structures are unavailable due to the difficulties to determine protein structures experimentally [14].

By comparison, sequence-based methods learn local patterns of metal ion-binding characteristics through sequence-derived features. For example, IonSeq [13] and TargetS [15] extract evolutionary conservative information, predicted secondary structure and ligand-specific binding propensity from sequence context using sliding-window strategy, and then employ support vector machine (SVM) to learn local binding patterns. Sequence-based approaches have a potentially wider range of applications since they require only readily available protein sequences, yet the lacks of tertiary structure information usually cause their limited performances.

In addition to the weaknesses described above, both structure-based and sequence-based approaches are mostly time-consuming owing to the usage of evolutionary sequence profile from multi-sequence alignment. For example, the profile generation using PSI-BLAST [16] requires about an hour for individual protein on single CPU, and the computational time is even growing due to the exponential growth of the sequence library. This has partly limited their large-scale applications in proteomes. Unsupervised pretraining with contextual language models has yielded ground-breaking improvements in natural language processing, which has recently been applied to protein sequence representation learning and has displayed highly promising results in downstream predictions including secondary structure, tertiary contact, mutational effect, and ontology-based protein function [17-19]. Such breakthroughs inspire us to develop a fast and accurate sequence-based metal ion-binding site predictor. Besides, current methods in this field learn different ion-binding patterns separately, ignoring the underlying relations between these similar ligands. Multi-task learning aims to improve predictive performance for related tasks (e.g. binding sites of different metal ions) by exploiting shared networks [20], which has been shown to benefit bioinformatics problems including predictions of inter-residue distances [21], cleavage sites [22], and binding sites [23, 24]. Therefore, it is promising to further advance metal ion-binding site prediction by effectively modeling the intrinsic relations between different ions through multi-task learning technique.

In this study, we present LMetalSite, a novel alignment-free sequence-based method for binding site predictions of the four most frequently seen metal ions (Zn^2+^, Ca^2+^, Mg^2+^ and Mn^2+^) in structures from PDB. LMetalSite leverages the recently published pretrained language model (ProtTrans [18]) to bypass slow database searches and generate informative sequence representations within a short time. Multi-task learning is adopted to further improve the predictive quality by compensating for the scarcity of training data and better modeling the intrinsic similarities between different metal ions. Concretely, we employ the well-acknowledged transformer model [25, 26] as shared networks to capture common binding mechanisms such as long-range dependencies in protein sequence, followed by four ion-specific multilayer perceptrons to learn the binding patterns of particular metal ions. LMetalSite was shown to surpass even the state-of-the-art structure-based methods by more than 19.7%, 14.4%, 36.8%, and 12.6% in AUPR on the four independent tests, respectively. Further analyses indicated that the self-attention modules are effective to learn the structural contexts of residues from protein sequence representation, which partly explains the superior performance of LMetalSite. In the future, our framework can be easily extended to sequence-based predictions of other functional sites, such as nucleic-acid-binding sites.

## 2. Materials and Methods

### 2.1 Datasets

To evaluate the performance of LMetalSite, we constructed four benchmark datasets for the four most frequently seen metal ion-binding proteins from the BioLiP database [27], including Zn^2+^, Ca^2+^, Mg^2+^ and Mn^2+^ ions. This database is a collection of biologically relevant protein-ligand complexes primarily from PDB. Concretely, we collected proteins that bind with these ions from BioLiP released on 29 December 2021. Only protein chains with resolutions of ≤ 3.0 Å and lengths of 50 to 1000 were kept. In these datasets, a binding residue was defined if the smallest atomic distance between the target residue and the ligand molecule is less than 0.5 Å plus the sum of the Van der Waal’s radius of the two nearest atoms. Then, we removed redundant proteins sharing sequence identity > 25% over 30% overlap within each dataset using CD-HIT [28]. Finally, each benchmark dataset was further split into a training set that contains proteins released before 1 January 2020, as well as an independent test set that contains proteins released from 1 January 2020 to 29 December 2021. Specifically, the training sets of Zn^2+^, Ca^2+^, Mg^2+^ and Mn^2+^ ions contain 1647, 1554, 1730 and 547 chains, respectively, and the corresponding independent test sets contain 211, 183, 235 and 57 chains, respectively. Details of the statistics of these benchmark datasets are given in **Table 1**, where the datasets are named by the amounts of the included proteins.

**Table 1.**
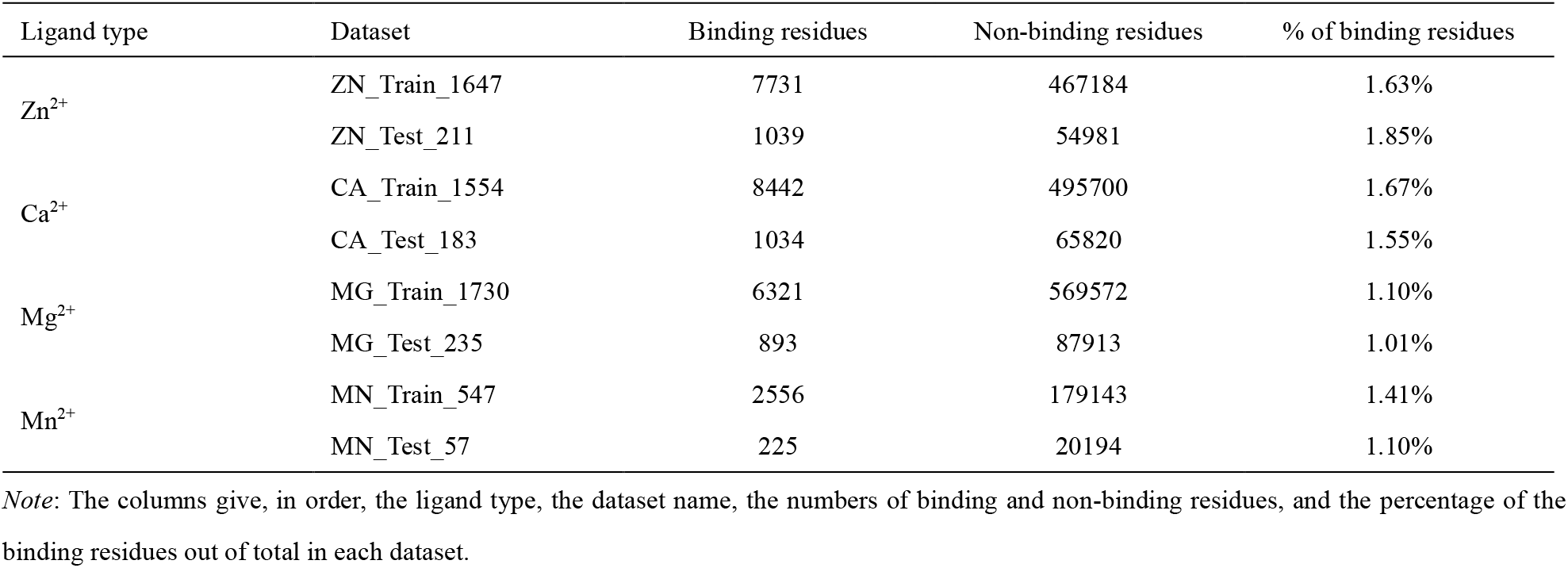
Statistics of the four metal ion benchmark datasets used in this study.

### 2.2 Protein sequence representation

LMetalSite solely employs pretrained language models to effectively generate informative sequence representation as input feature. Moreover, we have also tested other widely used features including evolutionary information and structural properties to demonstrate the superiority of the representation produced by pretrained language models (results are shown in **section 3.1**).

#### Language model representation

LMetalSite leverages the recent language model ProtT5-XL-U50 [18] (denoted as ProtTrans) for feature extraction, which is a transformer-based auto-encoder named T5 [29] pretrained on UniRef50 [30] in a self-supervised manner, essentially learning to predict masked amino acids. We extracted the output from the last layer of the encoder part of ProtTrans as sequence representation, which is a 1024-dimensional per-residue feature matrix. We have additionally investigated another similar language model, ESM-1b [17] (denoted as ESM), which was also pretrained on UniRef50 using transformer. Sequence representation by ESM is a 1280-dimensional per-residue feature matrix. Note that the inference costs of ProtTrans and ESM are really low, and the feature extraction process of our whole benchmark datasets (∼6000 sequences) using these pretrained models can be done within 10 minutes on an Nvidia GeForce RTX 3090 GPU.

#### Evolutionary information

Evolutionarily conserved residues may contain motifs related to important protein properties. Here, we also explored the widely used evolutionary features position-specific scoring matrix (PSSM) and hidden Markov models (HMM) profile. Concretely, PSSM was produced by running PSI-BLAST [16] to search the query sequence against UniRef90 [30] with three iterations and an E-value of 0.001. HMM profile was generated by running HHblits [31] to align the query sequence against UniClust30 [32] with default parameters. Each residue was encoded into a 20-dimensional vector in PSSM or HMM.

#### Structural properties

Three types of structural properties were extracted by DSSP [33] using native PDB structures: 1) 8-dimensional one-hot secondary structure profile. 2) Sine and cosine of the backbone torsion angles PHI and PSI. 3) Relative solvent accessibility, which is the normalized solvent accessible surface area (ASA) by the maximal ASA of the corresponding amino acid. This 13-dimensional structural feature is named DSSP hereinafter.

The feature values in the sequence representations from pretrained language models, PSSM, and HMM were all normalized to scores between 0 to 1 using Equation (1), where *v* is the original feature value, and *Min* and *Max* are the smallest and biggest values of this feature type observed in the training set.

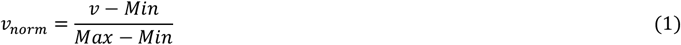

### 2.3 The architecture of LMetalSite

The overall architecture of the proposed framework LMetalSite is shown in **Figure 1**. First, the protein sequence is input to the pretrained language model to produce the sequence embedding, which is augmented by Gaussian noise to avoid overfitting. Then, the shared networks consisting of *N* transformer blocks are employed to capture the common binding-relevant characteristics such as long-range dependencies of the residues. Finally, four ion-specific multilayer perceptrons (MLPs) are adopted to learn the binding patterns of particular metal ions.

**Figure 1.**
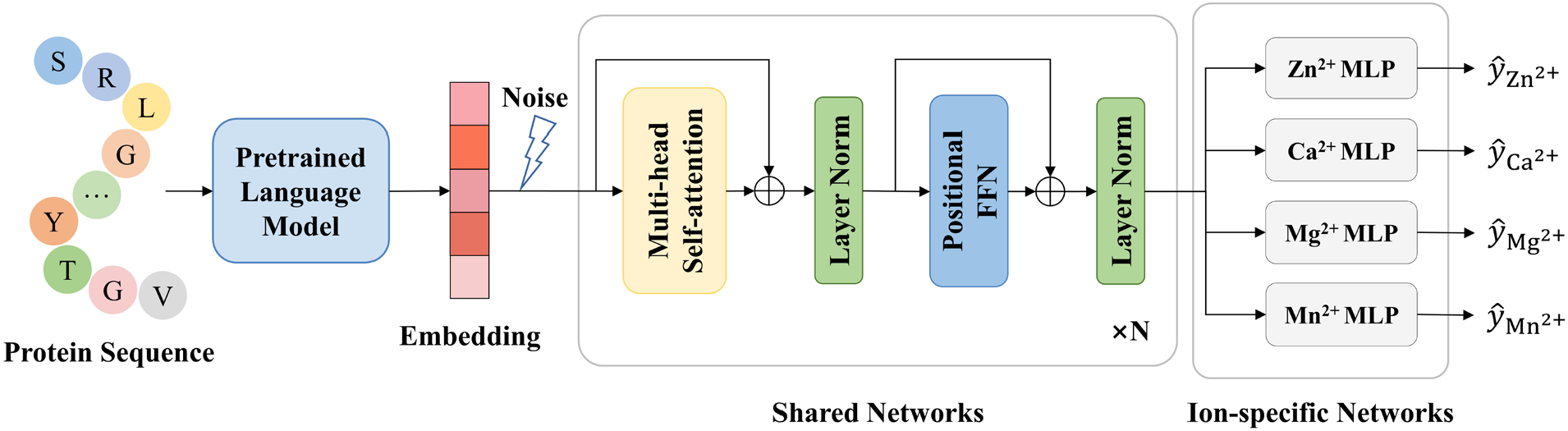
The overall architecture of the proposed framework LMetalSite. First, the protein sequence is input to the pretrained language model to produce the sequence embedding, which is augmented by Gaussian noise. Then, the shared transformer networks are employed to capture the common binding-relevant characteristics such as long-range dependencies. Finally, four ion-specific MLPs are adopted to learn the binding patterns of particular metal ions.

#### 2.3.1 Shared transformer networks

We stack multiple standard transformer encoder layers as shared networks to capture common binding characteristics of different metal ions. Each transformer layer consists of a multi-head self-attention module and a positional fully connected feed-forward network (FFN). A residual connection [34] is employed around each of the two sub-layers, followed by layer normalization [35]. Let 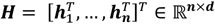 denote the input of the self-attention module, where *n* is the sequence length, *d* is the hidden dimension, and ***h***_*i*_ ∈ ℝ^1×*d*^ is the hidden representation of the *i*^th^ amino acid. The input of the *l*^th^ layer ***H***^(*l*)^ is projected by three matrices 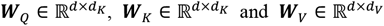 and 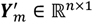 to the corresponding query, key and value representations ***Q, K, V***:

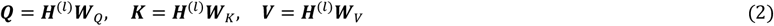

The self-attention is then calculated as:

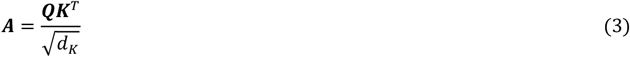

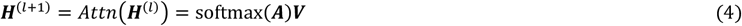

where ***A*** is a matrix capturing the similarities between queries and keys. To jointly attend to information from different representation subspaces at different positions, multi-head attention is used to linearly project the queries, keys and values *h* times, perform the attention function in parallel, and finally concatenate them together. In this study, *d*_*K*_ = *d*_*V*_ = *d* / *h*.

#### 2.3.2 Ion-specific multilayer perceptrons

The output of the last transformer layer is input to the ion-specific MLPs to predict the binding probabilities of particular metal ions for all *n* amino acid residues:

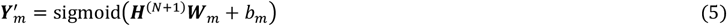

where ***H***^(*NN*+1)^ ∈ ℝ^*n*×*d*^ is the output of the *N*^th^ transformer layer; ***W***_*n*_ ∈ ℝ^*d*×1^ is the weight matrix for the specific metal ion *m*; *b*_*n*_ ∈ ℝ is the bias term for metal ion *m*, and ***Y′*** ∈ ℝ^*n*×1^ is the predictions of metal ion *m* for the *n* residues. The sigmoid function normalizes the output of the network into binding probabilities ranging from 0 to 1.

#### 2.3.3 Noise augmentation and multi-task training

Due to the limited training data and high dimension of the feature vectors, the sequence representation from the pretrained language model is augmented by Gaussian noise before being fed to the shared networks to avoid overfitting in the training steps:

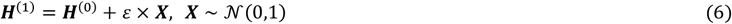

where ***H***^(0)^ is the sequence representation from the pretrained language model, ***X*** is a matrix with the same size as ***H***^(0)^ which is filled with random values from the standard normal distribution, and *ε* is a hyperparameter.

LMetalSite utilizes multi-task learning for simultaneous predictions of four metal ion-binding sites. Nevertheless, only the predictions of the corresponding known ion-binding types are used to calculate loss and perform backpropagation through masks in the training steps. That is, each protein is used to train the shared networks and the corresponding ion-specific network(s) of its known ion-binding type(s) without affecting other irrelevant MLPs.

### 2.4 Implementation details

We performed the 5-fold cross-validation on the training data, where the four training sets were mixed and split into five folds randomly, and then each time a model was trained on four folds and evaluated on the remaining fold. This process was repeated for five times and the performances on the five folds were averaged as the overall validation performance, which was used to choose the best feature combination and optimize all hyperparameters through grid search (**Supplementary Table S1**). In the test phase, all five trained models from the cross-validation were used to make predictions, which were averaged as the final predictions of LMetalSite.

Specifically, we employed a 2-layer shared transformer network with 64 hidden units and the following set of hyperparameters: *h* = 4, *ε* = 0.05 and batch size of 32. We utilized the Adam optimizer [36] with *β*_*1*_ = 0.9, *β*_*2*_ =0.99, weight decay of 10^−5^ and learning rate of 3 × 10^−4^ for model optimization on the binary cross-entropy loss. The dropout rate was set to 0.2 to avoid overfitting. We implemented the proposed model with Pytorch 1.7.1 [37]. Within each epoch, we randomly drew 30,000 samples from the training data with replacement to train our model. The training process lasted at most 30 epochs and we performed early-stopping with patience of 6 epochs based on the validation performance, which took about 45 minutes on an Nvidia GeForce RTX 3090 GPU. During the test phase, it took approximately 10 seconds to make predictions for one batch of proteins.

### 2.5 Evaluation metrics

Similar to the previous works [38, 39], we used recall (Rec), precision (Pre), F1-score (F1), Matthews correlation coefficient (MCC), area under the receiver operating characteristic curve (AUC), and area under the precision-recall curve (AUPR) to evaluate the predictive performance:

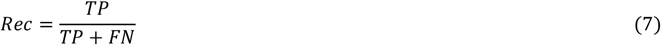

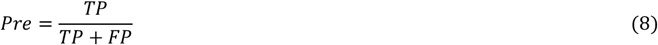

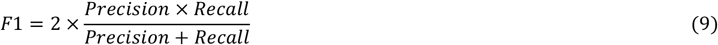

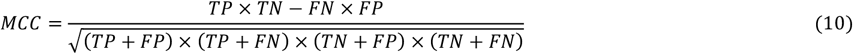

where true positives (TP) and true negatives (TN) denote the number of correctly predicted binding and non-binding residues, and false positives (FP) and false negatives (FN) denote the number of incorrectly predicted binding and non-binding residues, respectively. AUC and AUPR are independent of thresholds, thus reflecting the overall performance of a model. The other metrics were calculated using a threshold to convert the predicted binding probabilities to binary predictions, which was determined by maximizing MCC for the model. We adopted AUPR for hyperparameter selections as it is more sensitive and informative than AUC in imbalanced two-class classification tasks [40].

## 3 Results

### 3.1 Features from pretrained language models are informative for binding site detection

We evaluated LMetalSite by AUC and AUPR using the 5-fold cross-validation (CV) and independent test sets of Zn^2+^, Ca^2+^, Mg^2+^ and Mn^2+^ ions. As shown in **Table 2**, the LMetalSite model obtained average AUC values over the four metal ions of 0.913 and 0.928 on the 5-fold CV and independent tests, respectively; as well as average AUPR values of 0.540 and 0.559, respectively. The consistent performances on the CV and independent tests indicated the robustness of our model. Specifically, LMetalSite achieved AUPR of 0.803, 0.492, 0.316, and 0.625 on the four independent test sets.

**Table 2.** The performance of LMetalSite on the 5-fold CV and the four metal ion test sets using different features.

| Feature | CV | CV | Test | Test | Zn<sup>2+</sup> | Ca<sup>2+</sup> | Mg<sup>2+</sup> | Mn<sup>2+</sup> |
| --- | --- | --- | --- | --- | --- | --- | --- | --- |
|  | AUC* | AUPR* | AUC* | AUPR* | AUPR | AUPR | AUPR | AUPR |
| Evo | 0.853 | 0.374 | 0.870 | 0.392 | 0.694 | 0.239 | 0.173 | 0.462 |
| Evo+DSSP | 0.871 | 0.413 | 0.884 | 0.443 | 0.734 | 0.312 | 0.213 | 0.511 |
| ESM | 0.900 | 0.521 | 0.921 | 0.537 | 0.780 | 0.463 | 0.310 | 0.596 |
| ProtTrans | 0.913 | 0.540 | 0.928 | 0.559 | 0.803 | 0.492 | **0.316** | 0.625 |
| ProtTrans+ESM | 0.910 | 0.540 | 0.928 | 0.555 | 0.803 | 0.479 | 0.313 | 0.625 |
| ProtTrans+Evo+DSSP | **0.915** | **0.550** | **0.930** | **0.566** | **0.813** | **0.500** | 0.313 | **0.636** |

*Note:* \* denotes the average metrics of the four metal ions. Evo denotes the evolutionary features PSSM and HMM. Bold fonts indicate the best results.

To demonstrate the effect of sequence representation by the ProtTrans language model, we conducted feature ablation experiments to compare ProtTrans with another language model ESM and other widely used handcrafted features in this field. As shown in **Table 2**, when using the computationally efficient pretrained model ProtTrans or ESM to extract sequence representation, the model gained an average AUPR of 0.559 or 0.537 on the independent tests, respectively, higher than the one (0.443) by using evolutionary information (PSSM and HMM) and native structural features (DSSP). Note that ProtTrans performs slightly better than ESM, which might be ascribed to the different network architectures they adopted for pretraining. Moreover, combining ProtTrans and ESM is redundant and cannot attain any further improvement, suggesting that these two language models are similar. Additionally, further integrating evolutionary and structural features to ProtTrans only brings minor improvements on the test sets (less than 0.01 of AUPR on average). These results indicated that the ProtTrans language model may potentially capture the evolutionary and structural information of the protein.

### 3.2 The impact of multi-task learning

LMetalSite employs multi-task learning to capture the intrinsic similarities between different metal ions. To investigate the impact of multi-task technique, we changed the four ion-specific MLPs in LMetalSite into a single MLP, and then trained and tested on the four metal ion benchmark datasets separately with the same input sequence features. As shown in **Figure 2** and **Supplementary Table S2**, the removal of the multi-task strategy caused AUPR drops of 0.009, 0.007, 0.037, and 0.056 on the Zn^2+^, Ca^2+^, Mg^2+^ and Mn^2+^ test sets, respectively. As expected, the ion with the smallest training set (Mn^2+^) benefited the most from multi-task learning, since the transformer networks could be better trained with a larger dataset containing other types of metal ion-binding proteins. This suggested that different types of metal ions might potentially share common chemical mechanisms and binding patterns, and the predictions for one metal ion type could benefit from the binding information of other ion types. We have also shown that the Gaussian noise augmentation is effective in preventing overfitting, since its removal caused small but consistent performance drops in AUPR of 0.010, 0.028, 0.015, and 0.012 on the four test sets, respectively.

**Figure 2.**
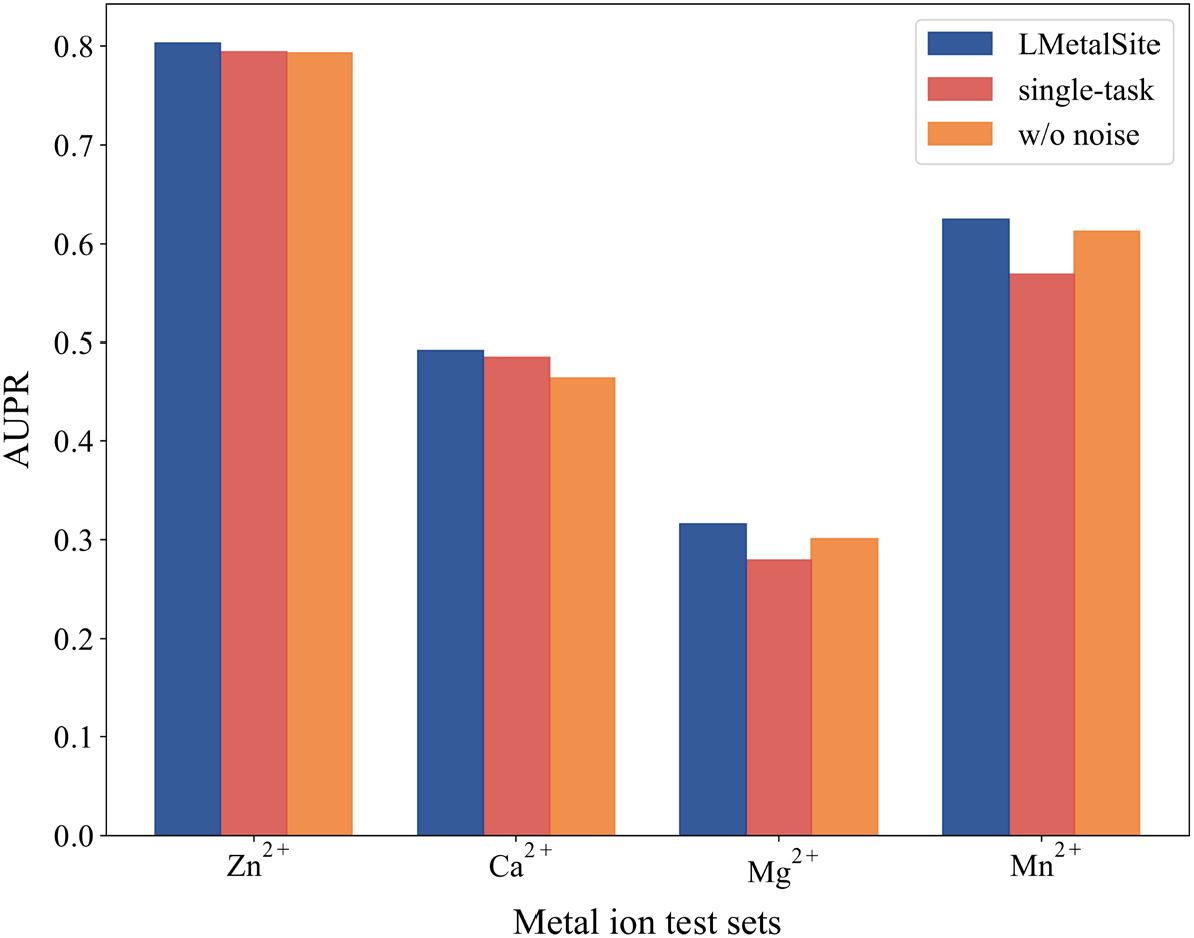
Ablation study on multi-task learning and noise augmentation in the four metal ion test sets.

### 3.3 Comparison with state-of-the-art methods

We compared LMetalSite with one sequence-based (TargetS) and four structure-based (MIB, IonCom, DELIA and GraphBind) predictors on the Ca^2+^, Mg^2+^ and Mn^2+^ test sets. Since binding site prediction of Zn^2+^ ion is not supported by DELIA and GraphBind, we only compared with the remaining three methods on the Zn^2+^ test set. As reported in **Table 3**, binding sites of Ca^2+^ and Mg^2+^ ions seem harder to distinguish, which may be due to the small differences in the binding propensities for Ca^2+^ and Mg^2+^ among different amino acids [15]. Howsoever, LMetalSite outperformed all other sequence-based and even structure-based methods in F1, MCC, AUC, and AUPR. Undoubtedly, LMetalSite substantially surpassed the sequence-based method TargetS by 35.4%, 201.8%, 113.5% and 94.1% in AUPR on Zn^2+^, Ca^2+^, Mg^2+^ and Mn^2+^ test sets, respectively. Interestingly, though our method is a sequence-based predictor, it outperformed the state-of-the-art structure-based method GraphBind by 14.4%, 36.8%, and 12.6% in AUPR on the Ca^2+^, Mg^2+^ and Mn^2+^ test sets, respectively. This is expected because the sequence representation from the pretrained language model used by LMetalSite is more informative and powerful than the handcrafted evolutionary and structural features employed by GraphBind. Another reason may be that the experimental structures also brought noises because the protein structures are flexible. In addition, the multi-task learning adopted by LMetalSite could further boost the predictive quality through better trained shared networks that capture common binding patterns among different ions. Our method also outperformed IonCom by 19.7% on the Zn^2+^ test set, likely because IonCom is a hybrid method and the newly resolved proteins didn’t always have high-quality templates. The improved performance of LMetalSite over other methods on the four metal ion test sets can be further illustrated by the precision-recall curves (**Figure 3**) and the ROC curves (**Supplementary Figure S1**), where the curves of LMetalSite are mostly located on the upper portion of the figures. Note that our method didn’t always have the highest recall and precision, because they are unbalanced measures strongly depending on thresholds. On the other hand, our method is also computationally efficient. Empirically, it takes less than 1 minute to make predictions for a protein with 200 residues using LMetalSite, but about 30 minutes using GraphBind or 6 hours using IonCom.

**Table 3.** Performance comparison of LMetalSite with state-of-the-art methods on the four metal ion test sets.

| Dataset | Method | Rec | Pre | F1 | MCC | AUC | AUPR |
| --- | --- | --- | --- | --- | --- | --- | --- |
| Zn<sup>2+</sup> | MIB | 0.744 | 0.219 | 0.339 | 0.385 | 0.935 | 0.394 |
|  | TargetS | 0.454 | 0.749 | 0.566 | 0.578 | 0.874 | 0.593 |
|  | IonCom | 0.852 | 0.137 | 0.236 | 0.317 | 0.937 | 0.671 |
|  | LMetalSite | 0.681 | 0.859 | **0.760** | **0.761** | **0.976** | **0.803** |
| Ca<sup>2+</sup> | MIB | 0.338 | 0.078 | 0.126 | 0.135 | 0.775 | 0.103 |
|  | TargetS | 0.121 | 0.490 | 0.194 | 0.238 | 0.776 | 0.163 |
|  | IonCom | 0.297 | 0.247 | 0.269 | 0.258 | 0.698 | 0.166 |
|  | DELIA | 0.172 | 0.633 | 0.271 | 0.325 | 0.785 | 0.248 |
|  | GraphBind | 0.371 | 0.623 | 0.465 | 0.475 | 0.888 | 0.430 |
|  | LMetalSite | 0.413 | 0.724 | **0.526** | **0.542** | **0.905** | **0.492** |
| $\text{Mg}^{2+}$ | MIB | 0.246 | 0.043 | 0.074 | 0.082 | 0.675 | 0.053 |
|  | TargetS | 0.118 | 0.491 | 0.190 | 0.237 | 0.724 | 0.148 |
|  | IonCom | 0.240 | 0.250 | 0.245 | 0.237 | 0.688 | 0.184 |
|  | DELIA | 0.129 | 0.650 | 0.215 | 0.287 | 0.744 | 0.198 |
|  | GraphBind | 0.273 | 0.414 | 0.329 | 0.331 | 0.776 | 0.231 |
|  | LMetalSite | 0.245 | 0.728 | **0.367** | **0.419** | **0.865** | **0.316** |
| $\text{Mn}^{2+}$ | MIB | 0.462 | 0.096 | 0.159 | 0.193 | 0.856 | 0.168 |
|  | TargetS | 0.271 | 0.496 | 0.351 | 0.362 | 0.864 | 0.322 |
|  | IonCom | 0.511 | 0.245 | 0.331 | 0.344 | 0.833 | 0.304 |
|  | DELIA | 0.502 | 0.665 | 0.572 | 0.574 | 0.902 | 0.489 |
|  | GraphBind | 0.427 | 0.706 | 0.532 | 0.545 | 0.930 | 0.555 |
|  | LMetalSite | 0.613 | 0.719 | **0.662** | **0.661** | **0.966** | **0.625** |

*Note:* The results of IonCom and GraphBind were obtained from their standalone programs, while the predictions by other competitive methods were generated from their web servers. Bold fonts indicate the best results.

**Figure 3.**
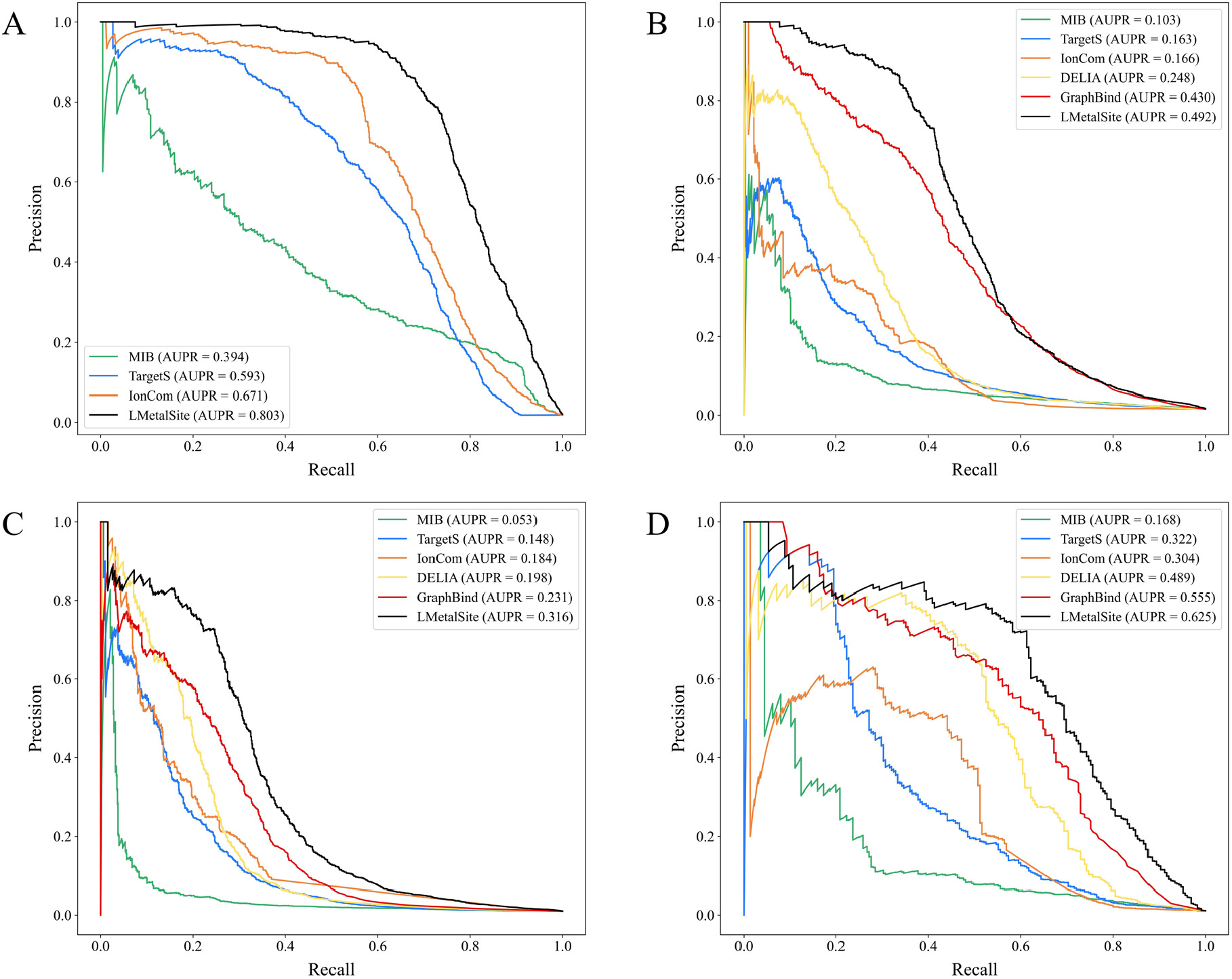
Precision-recall curves of LMetalSite and other methods on the Zn^2+^ (**A**), Ca^2+^ (**B**), Mg^2+^ (**C**) and Mn^2+^ (**D**) ion test sets.

### 3.4 Attention mechanism effectively captures protein structural information

To validate if our method can implicitly capture the structural information, we extracted the attention maps from the final layer of the shared transformer networks for all proteins in the four metal ion test sets, which were then averaged among the attention heads to obtain the final maps. We also collected the native PDB structures for these proteins to acquire the coordinate of the Cα atom in each residue, and then calculated the inter-residual Euclidean distance maps. Comparative study found that the average attention score of the residue pairs in contact (< 8 Å) is higher than those not in contact (7.02 × 10^−3^ vs 2.07 × 10^−3^). However, residues adjacent in sequence are inherently in contact spatially, so the non-local contacts are more meaningful to investigate, which are defined as the contacts between two residues that are ≥ 20 residues away in sequence positions, but < 8 Å in terms of Cα atomic distances. In addition, since the average sequence interval between residues not in contact is larger than those in non-local contact (194 and 93, respectively), we randomly sampled uncontacted residues to ensure the same sequence intervals as those in non-local contact to eliminate bias. Still, the average attention score of the residue pairs in non-local contact is significantly higher than the residue pairs not in contact after sampling (4.02 × 10^−3^ vs 2.67 × 10^−3^, *P*-value < 2.225 × 10^−308^) according to Mann–Whitney *U* test [41]. The distributions of the attention scores for the residue pairs in contact, in non-local contact, and not in contact are shown in **Figure 4**. These results suggested that the self-attention mechanism in LMetalSite effectively captures the protein structural information by spontaneously paying more attention to the spatial neighbors, which could facilitate better representation learning for the target residues as shown in our previous study [39].

**Figure 4.**
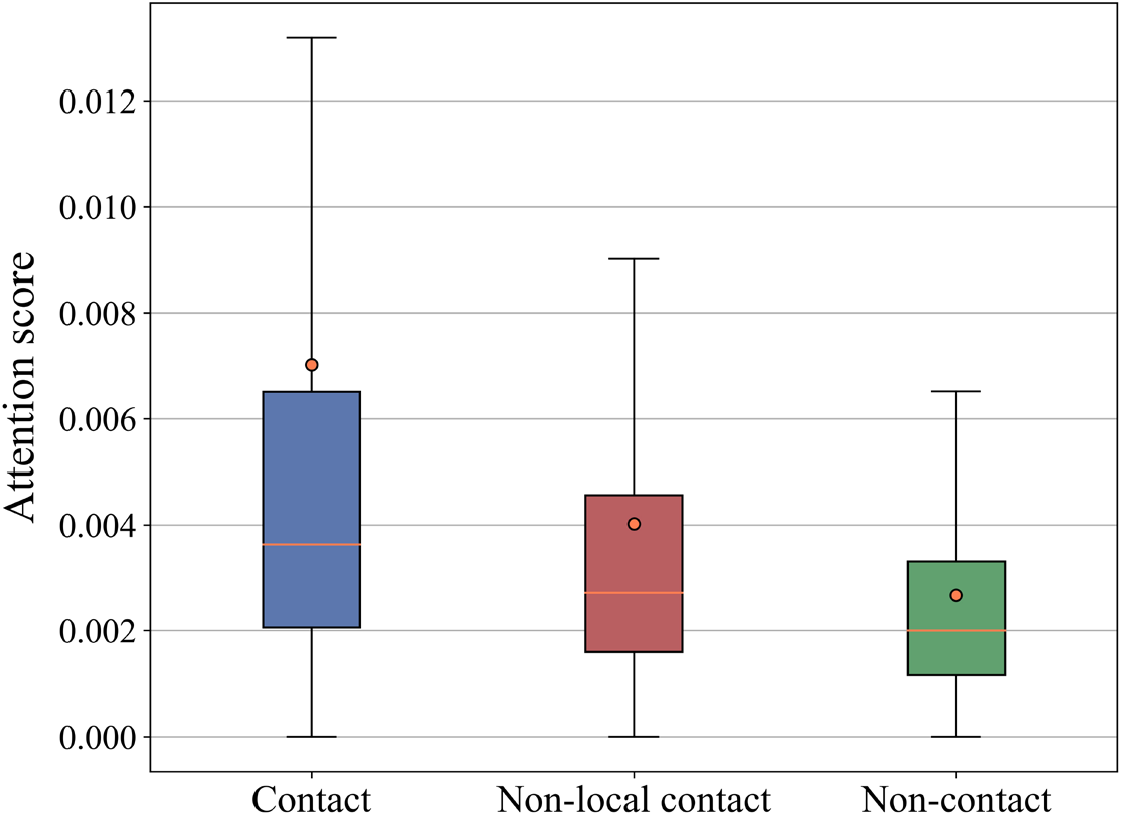
The distributions of the attention scores for the residue pairs in contact, in non-local contact, and not in contact. Each boxplot indicates the mean (orange dot), median (orange line), and quartiles with whiskers reaching up to 1.5 times the interquartile range. The outliers are not shown.

### 3.5 Case study

To visualize the superiority of our method, we selected the 40S ribosomal protein S29 (PDB ID: 6ZOJ, chain d) from ZN_Test_211 for illustration. **Figure 5A** shows its experimental contact map, where contacts were defined as the residue pairs with distances < 8 Å. We found that the non-local contacts generally match the residue pairs with high attention scores in the attention map from LMetalSite (**Figure 5B)**, suggesting that the attention module can partly capture the protein contact information. In this example, there are 4 Zn^2+^-binding residues over a total of 55 residues, and the prediction results of our method and the second-best method IonCom are shown in **Figure 5C** and **Figure 5D**, respectively. LMetalSite correctly predicted all binding residues with high confidences (probability scores > 0.99), and assigned non-binding residues with relatively low scores (< 0.24), leading to an MCC of 1.00. By comparison, IonCom predicted 16 binding residues in which 12 are false positives scattered spatially, leading to a lower MCC of 0.437.

**Figure 5.**
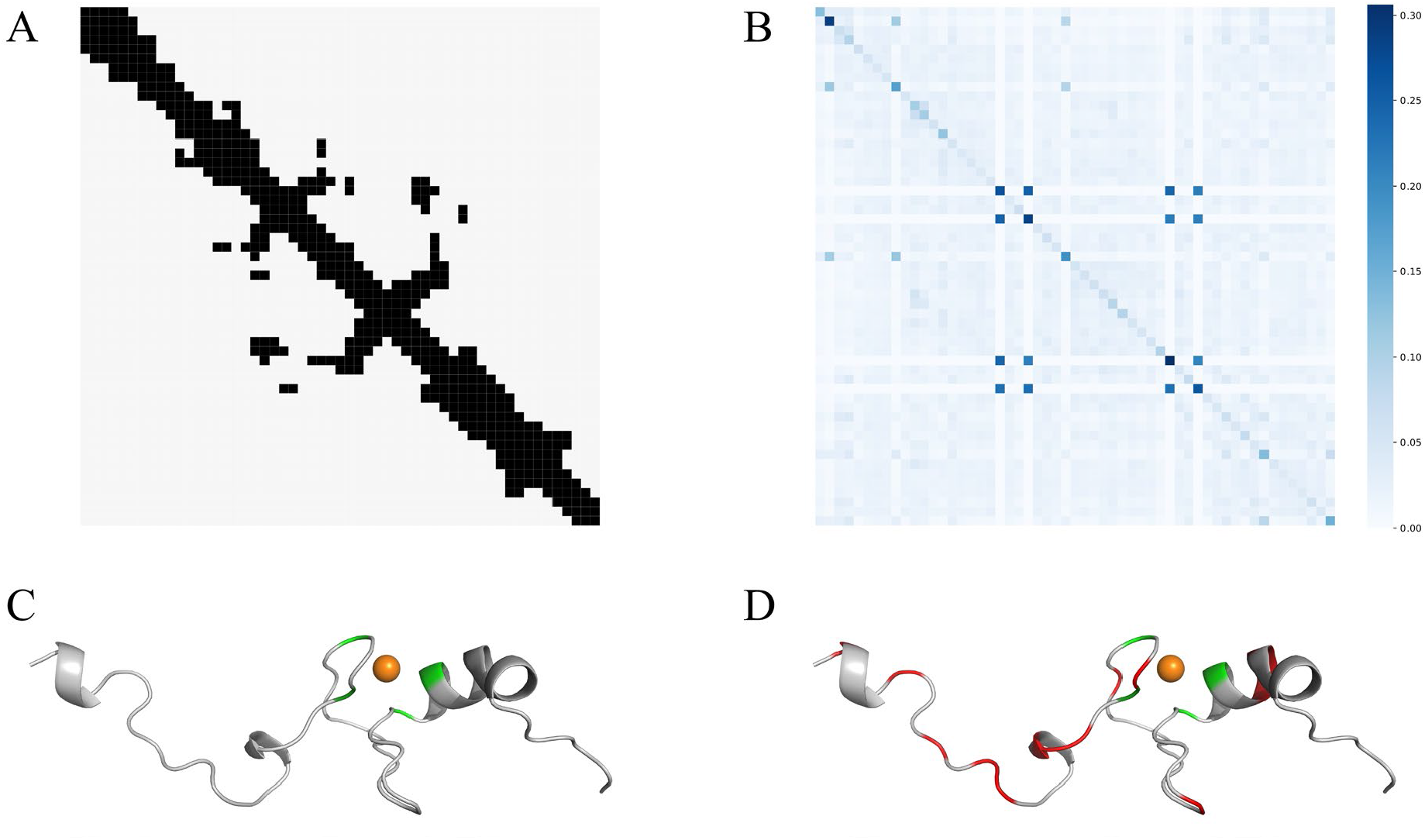
Visualization of one example (PDB ID: 6ZOJ, chain d) from ZN_Test_211. **A**. The experimental protein contact map. **B**. The attention map from LMetalSite. **C**. Predictions by LMetalSite. **D**. Predictions by IonCom. The orange spheres denote Zn^2+^ ions, and true positives and false positives are colored in green and red, respectively.

## 4 Discussion and Conclusion

Identifying metal ion-binding sites is crucial for understanding protein functions and designing novel drugs. Existing structure-based methods are not applicable to most proteins with unknown tertiary structures, while sequence-based methods are limited in predictive performance. Moreover, both structure-based and sequence-based approaches are mostly time-consuming owing to the usage of multi-sequence alignment. Trained with the computationally efficient and informative sequence representations from pretrained language model, LMetalSite achieved great performance (surpassing the best structure-based methods) using only protein sequences, which simultaneously solves the limitations of current structure-based and sequence-based methods. The multi-task learning technique adopted by LMetalSite is able to further improve the predictive quality, while all other competitive methods ignore the underlying relations between similar ions. In summary, the superiority of LMetalSite mainly benefits from two aspects: 1) the informative sequence representation from the pretrained language model potentially captures evolutionary and structural information; 2) multi-task learning is an effective algorithm to compensate for the scarcity of training data and better model the common binding mechanisms between different metal ions.

However, there is still room for further improvements on LMetalSite. Firstly, directly fine-tuning the pretrained language model on the binding site tasks may yield better performance than using it for feature exaction. Secondly, meta-learning [42, 43] could also be explored in the multi-task problems, which allows fast adaptation to unseen tasks with limited labels. Thirdly, although the pretrained language model may partly capture structural information such as secondary structure and solvent accessibility, and LMetalSite already exceeded the best structure-based methods, the binding site prediction could still benefit from known protein structures or high-quality predicted structures from AlphaFold2 [44] or RoseTTAFold [45]. For example, using distance maps to mask out spatially remote residues when calculating attention scores [39] or integrating pairwise geometric features between residues [46] may further enhance the predictive performance. Besides, spatial clustering of binding sites or constrained docking can be performed on the protein structures to additionally determine the coordinates of the binding locations.

In conclusion, this study proposed an alignment-free sequence-based framework LMetalSite for metal ion-binding site prediction, which employed the pretrained language model for sequence representation and multi-task learning to capture common binding patterns between different ions. LMetalSite showed preferable performance than other methods in comprehensive evaluations. We suggest that our method may provide useful information for biologists studying metal ion-binding patterns or pathogenic mechanisms of mutations, and chemists interested in targeted drug design. For example, the prediction can help narrow down potential binding sites for further wet experiment validation [47] or provide hypotheses and insights for the mechanisms of disease-causing gene mutations [48] as shown in other similar fields. The binding site prediction can also be used for druggability prediction [49] or de novo drug design [50-52]. In the future, we would further extend our method to predict various functional sites, including binding sites with proteins [38] and nucleic acids [39].

## Key points

- Existing structure-based methods for identifying metal ion-binding sites are not applicable to most proteins with unknown tertiary structures, while sequence-based methods are limited in predictive performance.
- LMetalSite is an alignment-free sequence-based method for metal ion-binding site prediction trained with informative sequence representation from the pretrained language model, which potentially captures evolutionary and structural information.
- LMetalSite employs multi-task learning to further improve the predictive quality by compensating for the scarcity of training data and better modeling the common binding patterns between different metal ions.
- LMetalSite showed preferable performance than state-of-the-art sequence-based and even structure-based methods in the four independent datasets of Zn^2+^, Ca^2+^, Mg^2+^ and Mn^2+^ ion-binding proteins.

## Supplementary Information

**Table S1.** The performance of LMetalSite on the 5-fold CV and the four metal ion test sets using different hyperparameters.

| Hyper-parameter | Value | CV AUPR* | Zn<sup>2+</sup> AUC | Ca<sup>2+</sup> AUC | Mg<sup>2+</sup> AUC | Mn<sup>2+</sup> AUC | Zn<sup>2+</sup> AUPR | Ca<sup>2+</sup> AUPR | Mg<sup>2+</sup> AUPR | Mn<sup>2+</sup> AUPR |
| --- | --- | --- | --- | --- | --- | --- | --- | --- | --- | --- |
| Layers | 1 | 0.536 | 0.975 | 0.905 | 0.861 | 0.967 | 0.800 | 0.493 | 0.305 | 0.619 |
|  | **2** | **0.540** | 0.976 | 0.905 | 0.865 | 0.966 | 0.803 | 0.492 | 0.316 | 0.625 |
|  | 3 | 0.539 | 0.975 | 0.904 | 0.861 | 0.963 | 0.799 | 0.492 | 0.304 | 0.622 |
|  | 4 | 0.538 | 0.973 | 0.904 | 0.871 | 0.961 | 0.802 | 0.494 | 0.306 | 0.616 |
|  | 5 | 0.539 | 0.974 | 0.904 | 0.867 | 0.960 | 0.800 | 0.491 | 0.301 | 0.627 |
| Hidden units | 32 | 0.537 | 0.975 | 0.904 | 0.867 | 0.964 | 0.799 | 0.487 | 0.304 | 0.607 |
|  | **64** | **0.540** | 0.976 | 0.905 | 0.865 | 0.966 | 0.803 | 0.492 | 0.316 | 0.625 |
|  | 128 | 0.537 | 0.975 | 0.902 | 0.858 | 0.965 | 0.797 | 0.487 | 0.309 | 0.617 |
| $h$ | 2 | 0.538 | 0.975 | 0.903 | 0.862 | 0.965 | 0.802 | 0.491 | 0.312 | 0.627 |
|  | **4** | **0.540** | 0.976 | 0.905 | 0.865 | 0.966 | 0.803 | 0.492 | 0.316 | 0.625 |
|  | 8 | 0.539 | 0.976 | 0.903 | 0.866 | 0.964 | 0.804 | 0.489 | 0.309 | 0.618 |
| $\varepsilon$ | 0 | 0.519 | 0.974 | 0.895 | 0.858 | 0.961 | 0.793 | 0.464 | 0.301 | 0.613 |
|  | 0.01 | 0.520 | 0.975 | 0.897 | 0.859 | 0.964 | 0.796 | 0.469 | 0.304 | 0.614 |
|  | **0.05** | **0.540** | 0.976 | 0.905 | 0.865 | 0.966 | 0.803 | 0.492 | 0.316 | 0.625 |
|  | 0.1 | 0.537 | 0.975 | 0.905 | 0.866 | 0.966 | 0.796 | 0.485 | 0.298 | 0.603 |
| Dropout | 0.1 | 0.534 | 0.975 | 0.905 | 0.866 | 0.964 | 0.806 | 0.487 | 0.306 | 0.627 |
|  | **0.2** | **0.540** | 0.976 | 0.905 | 0.865 | 0.966 | 0.803 | 0.492 | 0.316 | 0.625 |
|  | 0.3 | 0.539 | 0.974 | 0.904 | 0.865 | 0.967 | 0.799 | 0.486 | 0.305 | 0.618 |
| Learning rate | 1e-4 | 0.539 | 0.975 | 0.905 | 0.864 | 0.966 | 0.801 | 0.492 | 0.308 | 0.621 |
|  | **3e-4** | **0.540** | 0.976 | 0.905 | 0.865 | 0.966 | 0.803 | 0.492 | 0.316 | 0.625 |
|  | 5e-4 | 0.538 | 0.976 | 0.906 | 0.862 | 0.967 | 0.803 | 0.495 | 0.304 | 0.620 |

*Note:* For each row, the hyperparameters that are not shown are the same as the final LMetalSite model. CV AUPR\* denotes the average AUPR of the four metal ions on the 5-fold CV. Bold fonts indicate the best hyperparameters and the corresponding CV AUPR.

**Table S2.** Ablation study on multi-task learning and noise augmentation on the 5-fold CV and the four metal ion test sets.

| Method | CV | Zn<sup>2+</sup> | Ca<sup>2+</sup> | Mg<sup>2+</sup> | Mn<sup>2+</sup> | Zn<sup>2+</sup> | Ca<sup>2+</sup> | Mg<sup>2+</sup> | Mn<sup>2+</sup> |
| --- | --- | --- | --- | --- | --- | --- | --- | --- | --- |
|  | AUPR* | AUC | AUC | AUC | AUC | AUPR | AUPR | AUPR | AUPR |
| w/o noise | 0.519 | 0.974 | 0.895 | 0.858 | 0.961 | 0.793 | 0.464 | 0.301 | 0.613 |
| single-task | 0.512 | 0.975 | 0.895 | 0.857 | 0.960 | 0.794 | 0.485 | 0.279 | 0.569 |
| LMetalSite | **0.540** | **0.976** | **0.905** | **0.865** | **0.966** | **0.803** | **0.492** | **0.316** | **0.625** |

Note: CV AUPR\* denotes the average AUPR of the four metal ions on the 5-fold CV. The highest values are bolded.

**Figure S1.**
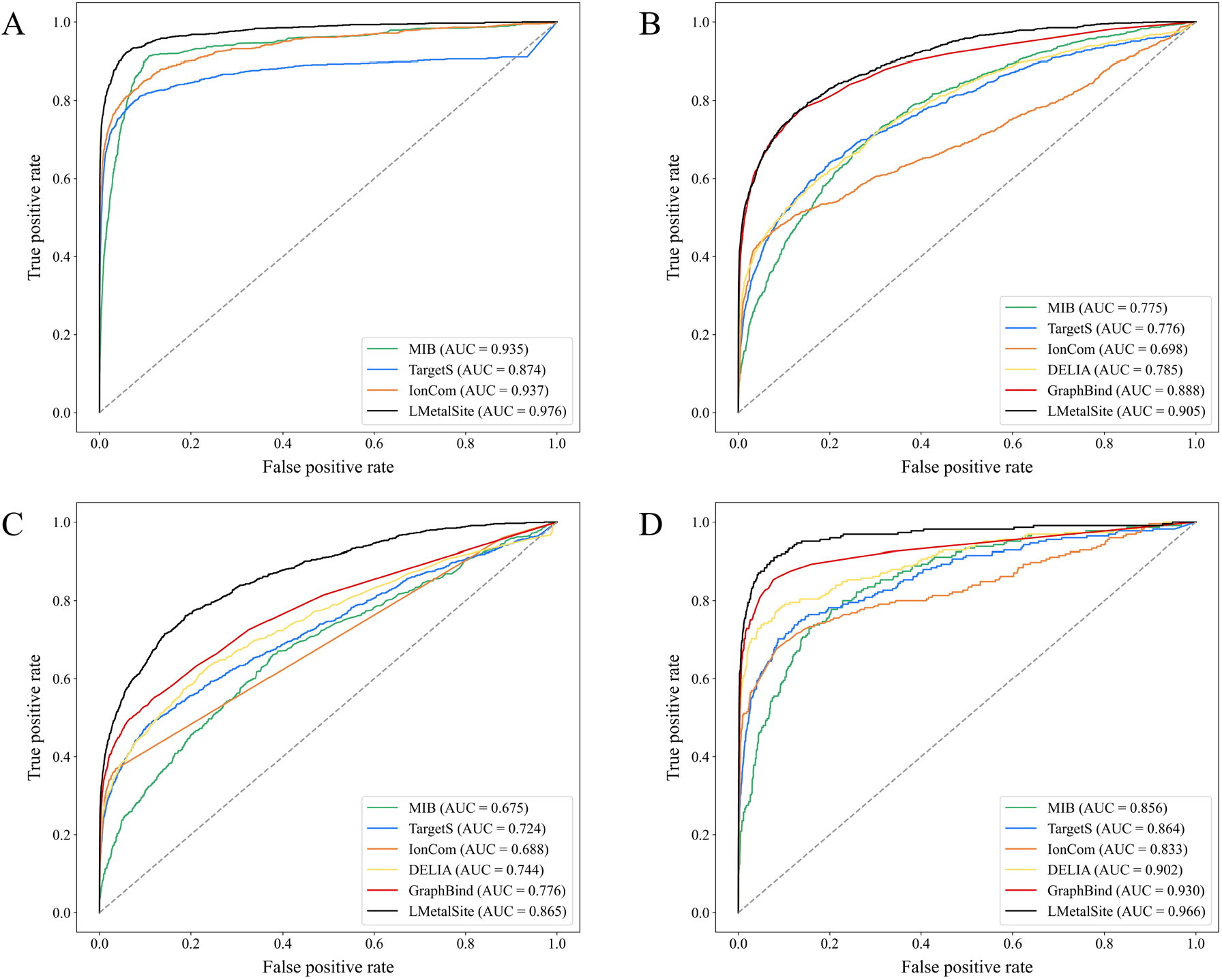
Receiver operating characteristic curves of LMetalSite and other methods on the Zn^2+^ (**A**), Ca^2+^ (**B**), Mg^2+^ (**C**) and Mn^2+^(**D**) ion test sets.

## References

1. Berman HM, Westbrook J, Feng Z et al. The protein data bank, Nucleic acids research 2000;28:235–242.

2. Putignano V, Rosato A, Banci L et al. MetalPDB in 2018: a database of metal sites in biological macromolecular structures, Nucleic acids research 2018;46:D459–D464.

3. Tainer JA, Roberts VA, Getzoff ED. Metal-binding sites in proteins, Current opinion in biotechnology 1991;2:582–591.

4. Andreini C, Bertini I, Rosato A. Metalloproteomes: a bioinformatic approach, Accounts of chemical research 2009;42:1471–1479.

5. Andreini C, Bertini I, Cavallaro G et al. Metal ions in biological catalysis: from enzyme databases to general principles, JBIC Journal of Biological Inorganic Chemistry 2008;13:1205–1218.

6. Berg JM. Zinc finger domains: hypotheses and current knowledge, Annual review of biophysics and biophysical chemistry 1990;19:405–421.

7. Yang J, Roy A, Zhang Y. Protein–ligand binding site recognition using complementary binding-specific substructure comparison and sequence profile alignment, Bioinformatics 2013;29:2588–2595.

8. Jensen MR, Petersen G, Lauritzen C et al. Metal binding sites in proteins: identification and characterization by paramagnetic NMR relaxation, Biochemistry 2005;44:11014–11023.

9. Reed GH, Poyner RR. Mn2+ as a probe of divalent metal ion binding and function in enzymes and other proteins, Metal ions in biological systems 2000:231–256.

10. Lin Y-F, Cheng C-W, Shih C-S et al. MIB: metal ion-binding site prediction and docking server, Journal of chemical information and modeling 2016;56:2287–2291.

11. Xia C-Q, Pan X, Shen H-B. Protein–ligand binding residue prediction enhancement through hybrid deep heterogeneous learning of sequence and structure data, Bioinformatics 2020;36:3018–3027.

12. Xia Y, Xia C-Q, Pan X et al. GraphBind: protein structural context embedded rules learned by hierarchical graph neural networks for recognizing nucleic-acid-binding residues, Nucleic acids research 2021;49:e51–e51.

13. Hu X, Dong Q, Yang J et al. Recognizing metal and acid radical ion-binding sites by integrating ab initio modeling with template-based transferals, Bioinformatics 2016;32:3260–3269.

14. Nagarajan R, Ahmad S, Michael Gromiha M. Novel approach for selecting the best predictor for identifying the binding sites in DNA binding proteins, Nucleic acids research 2013;41:7606–7614.

15. Yu D-J, Hu J, Yang J et al. Designing template-free predictor for targeting protein-ligand binding sites with classifier ensemble and spatial clustering, IEEE/ACM transactions on computational biology and bioinformatics 2013;10:994–1008.

16. Altschul SF, Madden TL, Schäffer AA et al. Gapped BLAST and PSI-BLAST: a new generation of protein database search programs, Nucleic acids research 1997;25:3389–3402.

17. Rives A, Meier J, Sercu T et al. Biological structure and function emerge from scaling unsupervised learning to 250 million protein sequences, Proceedings of the National Academy of Sciences 2021;118.

18. Elnaggar A, Heinzinger M, Dallago C et al. ProtTrans: Towards Cracking the Language of Lifes Code Through Self-Supervised Deep Learning and High Performance Computing, IEEE transactions on pattern analysis and machine intelligence 2021.

19. Unsal S, Atas H, Albayrak M et al. Learning functional properties of proteins with language models, Nature Machine Intelligence 2022;4:227–245.

20. Zhang Y, Yang Q. An overview of multi-task learning, National Science Review 2018;5:30–43.

21. Wu T, Guo Z, Hou J et al. DeepDist: real-value inter-residue distance prediction with deep residual convolutional network, BMC bioinformatics 2021;22:1–17.

22. Singh D, Sisodia DS, Singh P. Compositional framework for multitask learning in the identification of cleavage sites of HIV-1 protease, Journal of Biomedical Informatics 2020;102:103376.

23. Sun Z, Zheng S, Zhao H et al. To improve the predictions of binding residues with DNA, RNA, carbohydrate, and peptide via multi-task deep neural networks, IEEE/ACM transactions on computational biology and bioinformatics 2021.

24. Zhang F, Zhao B, Shi W et al. DeepDISOBind: accurate prediction of RNA-, DNA-and protein-binding intrinsically disordered residues with deep multi-task learning, Briefings in Bioinformatics 2022;23:bbab521.

25. Vaswani A, Shazeer N, Parmar N et al. Attention is all you need. In: Advances in neural information processing systems. 2017, p. 5998–6008.

26. Zheng S, Rao J, Zhang Z et al. Predicting retrosynthetic reactions using self-corrected transformer neural networks, Journal of chemical information and modeling 2019;60:47–55.

27. Yang J, Roy A, Zhang Y. BioLiP: a semi-manually curated database for biologically relevant ligand–protein interactions, Nucleic acids research 2012;41:D1096–D1103.

28. Fu L, Niu B, Zhu Z et al. CD-HIT: accelerated for clustering the next-generation sequencing data, Bioinformatics 2012;28:3150–3152.

29. Raffel C, Shazeer N, Roberts A et al. Exploring the Limits of Transfer Learning with a Unified Text-to-Text Transformer, Journal of machine learning research 2020;21:1–67.

30. Suzek BE, Huang H, McGarvey P et al. UniRef: comprehensive and non-redundant UniProt reference clusters, Bioinformatics 2007;23:1282–1288.

31. Remmert M, Biegert A, Hauser A et al. HHblits: lightning-fast iterative protein sequence searching by HMM-HMM alignment, Nature methods 2012;9:173–175.

32. Mirdita M, von den Driesch L, Galiez C et al. Uniclust databases of clustered and deeply annotated protein sequences and alignments, Nucleic acids research 2017;45:D170–D176.

33. Kabsch W, Sander C. Dictionary of protein secondary structure: pattern recognition of hydrogen-bonded and geometrical features, Biopolymers: Original Research on Biomolecules 1983;22:2577–2637.

34. He K, Zhang X, Ren S et al. Deep residual learning for image recognition. In: Proceedings of the IEEE conference on computer vision and pattern recognition. 2016, pp. 770–778.

35. Ba JL, Kiros JR, Hinton GE. Layer Normalization, stat 2016;1050:21.

36. Kingma DP, Ba J. Adam: A Method for Stochastic Optimization. In: 3rd International Conference on Learning Representations (Poster). 2015.

37. Paszke A, Gross S, Massa F et al. Pytorch: An imperative style, high-performance deep learning library, Advances in neural information processing systems 2019;32:8026–8037.

38. Yuan Q, Chen J, Zhao H et al. Structure-aware protein–protein interaction site prediction using deep graph convolutional network, Bioinformatics 2022;38:125–132.

39. Yuan Q, Chen S, Rao J et al. AlphaFold2-aware protein–DNA binding site prediction using graph transformer, Briefings in Bioinformatics 2022.

40. Saito T, Rehmsmeier M. The precision-recall plot is more informative than the ROC plot when evaluating binary classifiers on imbalanced datasets, PloS one 2015;10:e0118432.

41. Mann HB, Whitney DR. On a test of whether one of two random variables is stochastically larger than the other, The annals of mathematical statistics 1947:50–60.

42. Finn C, Abbeel P, Levine S. Model-agnostic meta-learning for fast adaptation of deep networks. In: International conference on machine learning. 2017, pp. 1126–1135. PMLR.

43. Wang J, Zheng S, Chen J et al. Meta learning for low-resource molecular optimization, Journal of Chemical Information and Modeling 2021;61:1627–1636.

44. Jumper J, Evans R, Pritzel A et al. Highly accurate protein structure prediction with AlphaFold, Nature 2021:1–11.

45. Baek M, DiMaio F, Anishchenko I et al. Accurate prediction of protein structures and interactions using a three-track neural network, Science 2021;373:871–876.

46. Ingraham J, Garg V, Barzilay R et al. Generative Models for Graph-Based Protein Design, Advances in neural information processing systems 2019;32:15820–15831.

47. Wang S, Liang K, Hu Q et al. JAK2-binding long noncoding RNA promotes breast cancer brain metastasis, The Journal of clinical investigation 2017;127:4498–4515.

48. Kumar R, Corbett MA, Van Bon BW et al. THOC2 mutations implicate mRNA-export pathway in X-linked intellectual disability, The American Journal of Human Genetics 2015;97:302–310.

49. Schmidtke P, Barril X. Understanding and predicting druggability. A high-throughput method for detection of drug binding sites, Journal of medicinal chemistry 2010;53:5858–5867.

50. Xu M, Ran T, Chen H. De novo molecule design through the molecular generative model conditioned by 3D information of protein binding sites, Journal of Chemical Information and Modeling 2021;61:3240–3254.

51. Zheng S, Li Y, Chen S et al. Predicting drug–protein interaction using quasi-visual question answering system, Nature Machine Intelligence 2020;2:134–140.

52. Wang P, Zheng S, Jiang Y et al. Structure-Aware Multimodal Deep Learning for Drug–Protein Interaction Prediction, Journal of chemical information and modeling 2022;62:1308–1317.

